# Repellency effects of nanoliposomal gels bearing Clove (S*yzygium aromaticum*) or Tea tree (*Melaleuca alternifolia*) essential oils against the main malaria vector, *Anopheles stephensi*

**DOI:** 10.64898/2026.07.07.736909

**Authors:** Raziyeh Shahheidari, Mohammad Djaefar Moemenbellah-Fard, Mahmoud Osanloo, Azim Paksa, Amir Hossein Roozitalab, Mohsen Fakhraei, Elham Zarenezhad

**Affiliations:** Student Research Committee, Department of Biology and Control of Disease Vectors, School of Health, Shiraz University of Medical Sciences, Shiraz, Iran; Research Center for Health Sciences, Institute of Health, Department of Biology and Control of Disease Vectors, School of Health, Shiraz University of Medical Sciences, Shiraz, Iran; Department of Medical Nanotechnology, School of Advanced Technologies in Medicine, Fasa University of Medical Sciences, Fasa, Iran; Noncommunicable Diseases Research Center, Fasa University of Medical Sciences, Fasa, Iran

**Author notes:** **<u>Corresponding Authors</u>:** Mohammad Djaefar **Moemenbellah-Fard** (PhD), Mahmoud Osanloo (PhD).

**Keywords:** *Anopheles stephensi*, Repellency, Mosquito, Nanoliposome, Volatile oil, Tea, Clove

## Abstract

**Background:** The development of safe and effective plant-based repellents is crucial to control malaria transmission, particularly given the spread of insecticide resistance in major vectors like *Anopheles stephensi*. Essential oils (EOs) are promising candidates, yet their high volatility and hydrophobicity limit their efficacy. This study aimed to design and evaluate nanoliposomal gels containing *Syzygium aromaticum* (clove) or *Melaleuca alternifolia* (tea tree) EOs to enhance their repellent durability against *An. stephensi*.

**Methods:** The chemical profiles of the EOs were determined via Gas Chromatography-Mass Spectrometry (GC-MS). Nanoliposomes bearing 3% of each EO were made ready with the ethanol injection method, and incorporated into a carboxymethyl cellulose (CMC) gel. Formulations were characterized for particle size, zeta potential, viscosity, and chemical interactions (FTIR). Repellent efficacy was evaluated using the arm-in-cage method, recording the complete protection time (CPT) for nanoliposomal gels (LipoGel 3%), in comparison with nonformulated EOs and the gold-standard repellent, DEET (40%).

**Results:** GC-MS analysis identified eugenol (79.51%) and terpinen-4-ol (73.53%) as the major constituents of clove and tea tree EOs, respectively. Nanoliposomes exhibited sizes of 82.3 ± 3 nm (clove) and 102 ± 4 nm (tea tree), with narrow size distributions. The clove LipoGel demonstrated a significantly enhanced CPT (341 ± 17 min), which was statistically comparable to 40% DEET (351 ± 16 min, *P*>0.05). In contrast, the nonformulated EOs resulted in only 45 min of protection, highlighting the critical role of the nanocarrier system.

**Conclusion:** The nanoliposomal gel formulation, particularly containing clove EO, represents a potent and safe botanical alternative to conventional synthetic repellents. This approach offers a promising strategy for integrated vector management, warranting further field-based investigations.

## 1. Introduction

Despite decades of global progress, malaria continues to adapt faster than many of the tools designed to eliminate it. One of the most striking examples of this adaptability is the emergence, dissemination, and proliferation of *Anopheles stephensi* in urban and man-made environments. It has even been recognized recently as an invasive species on the African continent (1).

The cause of this disease is a protozoan parasite transmitted between people by a group of blood-sucking mosquitoes (2). Children less than 5 years old, pregnant women, travelers, and individuals with AIDS are at risk of severe infection (3). Between 2000 and 2019, the estimated annual number of malaria cases remained stable, ranging from 227 to 248 million across the 108 malaria-endemic countries in 2000. Since 2020, the estimated number of malaria cases has steadily increased, with most of this rise occurring in countries within the WHO African Region (89.7%) and the WHO Eastern Mediterranean Region (15.5%) (4).

According to the World Health Organization, 263 million in 2023 and 282 million malaria cases in 2024 were reported. The number of deaths was also estimated at 597,000 in 2023 and 610,000 in 2024 (4, 5). Although the mortality rate for children under 5 has been declining, it still accounted for about 73% of malaria deaths worldwide (5).

In Iran in 2022, malaria cases increased significantly (1,439 local cases) due to the presence of expatriates, heavy rains, increased malaria cases in neighboring Pakistan, and poor disease detection in the southeastern region of the country, especially Sistan and Baluchestan province. The COVID-19 pandemic has also been identified as one of the reasons for the spread of this disease. In addition, warmer and wetter weather is predicted for this province, so there is a possibility of an annual outbreak in this region (6).

Recent studies have shown that due to climate change, the spread of *An. stephensi* is increasing in other regions of the world, particularly in Africa (7). The diversity of habitats of this species (*e.g.* water reservoirs, tree holes, and water streams), and its adaptation to different environments, have made it difficult to control (8). On the other hand, excessive use of larvicides and adulticides has caused resistance in *An. stephensi*. Therefore, the use of existing chemical larvicides and insecticides to control malaria vectors, especially *An. stephensi*, is not to the best interests of humans and the environment (9, 10).

Repellents play a crucial role in protecting people against arthropods since they can be used anytime and anywhere. They help prevent vector-borne diseases by reducing contact between humans and vectors (11). The most effective and widely used standard insect repellent is meta-diethyl-N,N-toluamide (DEET), which has been commercially available since the 1950s. This compound is effective for repelling mosquitoes, ticks, and other biting insects. Although DEET is highly effective, some insects are not affected by its application. A genetic foundation for this has been demonstrated in *Aedes aegypti* and fruit flies (12). On the other hand, toxicity related to DEET has been reported. This product can irritate mucous membranes and may also cause encephalopathy in children (11). Chronic DEET abuse has also been reported to affect nerve cells, cause abnormalities and possible neurobehavioral effects in adult rats (13).

The use of herbal-based repellents can thus be a good alternative to chemical pesticides. The benefits of plant-based formulations include their novelty and reduction in the development of resistance in insects (14). Many botanicals have shown a wide range of anti-nutritional, ovipository, inhibitory, growth-regulatory, repellency, larvicidal, and insecticidal activities (15, 16). Since botanical insecticides are generally non-allergenic to humans and act specifically, they can be considered as a good alternative to chemical pesticides (17–19).

Clove (*Syzygium aromaticum*) is a well-known aromatic herb with medicinal value, traditionally classified within the Myrtaceae family. It is primarily native to Indonesia, India, and Madagascar (20). Clove has been extensively studied for its diverse properties and applications. Research has demonstrated its effectiveness against various pathogenic parasites and microorganisms, including bacteria, *Plasmodium*, *Babesia*, and Herpes simplex virus, showing promising therapeutic effects (21). Various reports have also noted the repellency of clove essential oil (EO) against important medical vectors, including *An. stephensi*, *Culex pipiens* and *Ae. aegypti* (22–24).

The tea tree, *Melaleuca alternifolia*, is an aromatic plant of the family Myrtaceae and native to Australia (25). The EO extracted from its leaves has long been valued for its diverse biological properties. Tea tree oil is full of active compounds, particularly terpinen-4-ol, which plays a major role in its antimicrobial and repellent effects (26, 27). Due to these properties and its relative safety for humans, this EO is considered an attractive and natural option to reduce human contact with mosquitoes by disrupting their host-seeking behavior and thereby diminishing blood-feeding activity (28). The antimicrobial, antifungal, and anti-inflammatory properties of tea tree EO have been demonstrated in various studies. Consequently, it has been used to treat lice and *Demodex* infestations (29).

Plant EOs contain volatile compounds that are sensitive to some environmental conditions (oxidation, light, and air) and are hydrophobic. To overcome the hydrophobicity of volatile oils, liposomal formulations are currently used, in which oil droplets are encapsulated within a lipid shell (11). One suitable strategy to enhance the stability and efficacy of herbal EOs is to formulate them as liposomal nanoparticles. Liposomes are microscopic vesicles resembling cell membranes that can carry various substances either on their surfaces or interiors. Liposomes enhance clinical efficacy and drug delivery by improving drug loading capacity and providing sustained release of active compounds (11). Nanoliposomes possess several advantageous attributes, including biodegradability, amphiphilicity, non-immunogenicity, and low intrinsic toxicity (15). In addition, nanogels can be produced by adding substances to nanoliposomes. Nanogels, with their properties such as biocompatibility and biodegradability, high loading capacity, and appropriate viscosity, are suitable compounds for topical use as insect repellents (30).

Although clove EO has previously demonstrated repellent activity against *Anopheles* species, its protection time remains limited in conventional formulations (31). To enhance its persistence, the present study encapsulated clove oil in a nanoliposomal gel system. Additionally, tea tree oil, recognized for its low toxicity and promising bioactivity, but underexplored as a mosquito repellent, was also formulated as a nanoliposomal gel to evaluate its protective potential. This research aimed to determine whether nanoliposomal encapsulation can enhance overall repellency and extend the complete protection time (CPT) against *An. stephensi*. All formulations were compared to the standard DEET (40% w/v) repellent.

## 2. Materials and Methods

### 2.1 Materials

Essential oils (EOs) of *Syzygium aromaticum* (SAEO) and *Melaleuca alternifolia* (MAEO) were bought from Green Plant of Life Pharmaceutical Company, Iran. Carboxymethyl cellulose (CMC), absolute ethanol, wool fat cholesterol, and egg yolk lecithin were supplied from Merck Chemicals, Germany. DEET 40% was obtained from Reyhan Naqsh-e Jahan Pharmaceutical Company, Iran.

### 2.2 Mosquito rearing

Fasting female *An. stephensi* mosquitoes (5-7 day-old) were handled for the repellency experiments. These mosquitoes were reared in the insectarium of the Department of Biology and Control of Disease Vectors, Shiraz University of Medical Sciences, Shiraz, Iran, under the controlled conditions of 27±2°C, 70% relative humidity, and 12-h L/D cycle. Mosquitoes were unexposed to any insecticides. Adult female mosquitoes were blood-fed percutaneously on laboratory mice in accordance with the national guidelines (Ethical code number: IR.SUMS.SCHEANUT.REC.1404.02), and the international ethical guidelines of Helsinki. In addition to three blood meals per week, mosquitoes were also fed on a 10% sugar solution. Forty-eight hours after the blood meal, a porcelain bowl filled to approximately two-thirds of its capacity with clean water was placed inside the mosquito cage to facilitate oviposition. The deposited eggs were subsequently collected and transferred to larval rearing trays, where the larvae were nourished with fish food. After the required number of adult mosquitoes emerged, they were utilized for the repellency bioassays.

### 2.3 Characterization of constituents of SAEO and MAEO with GC-MS

Gas Chromatography–Mass Spectrometry (GC–MS) was employed to determine the chemical profiles of SAEO and MAEO. These analyses were performed using an Agilent 6890 GC system coupled to a 5973 mass selective detector (Agilent Technologies, Santa Clara, CA, USA). Separation of the EO constituents was achieved on a BPX5 fused-silica capillary column (30 m×0.25 mm, 0.25 μm film thickness). One µL of each oil sample, diluted in n-hexane, was filled into the instrument. The oven temperature program began at 50°C for 5 min, followed by an increase of 3°C/min up to 240°C, and then at 15°C/min to 300°C, where it was held for an additional 3 min, resulting in a total run time of 75 min. Injector and detector temperatures were set at 250°C and 230°C, respectively. The GC was operated with a split flow of 35 mL/min and a column flow of 1 mL/min. High-purity helium (99.99%) served as the carrier gas at a flow rate of 0.5 mL/min. Mass spectra were acquired at 70 eV using electron ionization with the ion source maintained at 220 °C, and data were recorded over an m/z range of 40–500.

Data processing was conducted with ChemStation software. Identification of the volatile components was carried out by comparing their retention indices, calculated using a homologous n-alkane series, with reference values reported in the literature and following the protocol explained in our previous publication (32).

### 2.4 Preparation of nonformulated EO solutions

To compare nonformulated essential oils with nanogels, a 3% (w/v) solution of clove EO (*Syzygium aromaticum*) or tea tree EO (*Melaleuca alternifolia*) was prepared. Each EO was dissolved in absolute ethanol as the solvent to obtain a final concentration of 3%. Ethanol was selected as the solvent to match the base solvent used in the nanoformulation system and to ensure complete dissolution of the EOs. Fresh solutions were prepared immediately before the bioassays.

### 2.5 Preparation and Characterization of Nanoliposomal Gels with the SAEO or MAEO

The initial nanoliposomes were prepared by ethanol injection method. First, a stock solution of lecithin and cholesterol (15% and 2.5%) was prepared using absolute ethanol (for 24 hours at room temperature, 2000 rpm). First, one mL of the oil phase was poured into the vial. Then, 3% w/v of EO was added to it. Subsequently, 0.5% w/v Tween 80 surfactants were added, and mixed for 3 minutes. Then, 4 mL of distilled water was added dropwise and mixed for 40 minutes.

To prepare the gel for repellency, 3.5% w/v of carboxymethyl cellulose (CMC) was added to the prepared nanoliposome containing 3% EOs and mixed overnight. The gel was stored at two temperatures (4°C and 26°C) for six months and was monitored for sedimentation, creaming, and biphasic separation.

The mean particle size of the nanoliposomes was measured using dynamic light scattering (DLS) with a K-One Nano particle size analyzer (K-One Nano Ltd., Korea). The particle size distribution (span) was calculated using the formula (d90 − d10)/d50, where d10, d50, and d90 indicate the particle diameters below which 10%, 50%, and 90% of the particles are found, respectively. The surface charge (zeta potential) of the nanoliposomes was measured using a Horiba DLS analyzer.

Attenuated total reflection-Fourier transform infrared (ATR-FTIR) spectroscopy was performed to verify the successful incorporation of the essential oils (EOs) into the nanogel formulation. Prior to analysis, the EO-loaded nanogels and the blank nanogel (without EO) were centrifuged at 12,000 × g for 60 min at 4°C. The resulting pellets were dried at room temperature for three days to minimize residual moisture before spectral analysis using a Bruker Tensor II FTIR spectrometer (USA). Infrared spectra were acquired over the wavenumber range of 400–4000 cm□^1^.

Viscometry analysis was used to measure the viscosity of the prepared gels. Viscosity was investigated at different shear rates (33).

### 2.6 Repellent Assays

Repellency measurements were performed under insectarium conditions according to the World Health Organization (WHO) guidelines for the arm-in-cage method, with minor modifications (34). Wooden cubic cages measuring 40×40×40 cm were used, surrounded by fine mesh netting equipped with a cloth sleeve at the front. The volunteer was a nonsmoker and avoided using scented products or consuming foods bearing odorous or mosquito-repellent compounds. Prior to testing, the forearm was washed with unscented soap, disinfected with 70% ethanol, and allowed to air dry. An area of 85 cm² of bare forearm skin was exposed for the assay, while the remaining arm was covered from the fingers to the elbow (34).

First, to verify the mosquitoes’ tendency to bloodfeed on human skin, the person’s hand was placed for three minutes without using any substance in a cage harboring 200 female mosquitoes that had been deprived of sugar water for 14 hours before the start of the test. To achieve the complete protection time (CPT) of the prepared materials, 1 ml of each sample (clove or tea tree EO solutions at 3% concentration, lipogels with either clove or tea tree EOs at 3% concentrations, blank lipogel, or 40% DEET) were applied to the volunteer’s hand. The volunteer’s forearm was then placed in the cage for three minutes, and this was repeated after each 30-minute period. The CPT was defined as the time interval between topical application of the sample and the occurrence of two landings in a 3-minute test or one landing in two consecutive 3-minute tests. All experiments were conducted in triplicate, and the complete protection times (CPTs) were expressed as mean ± standard deviation (SD). Statistical comparisons among the final values of all treatment groups were performed using one-way analysis of variance (ANOVA) in SPSS software (version 27).

## 3. Results

There were 10 and 19 different components identified in SAEO and MAEO using GC-MS analysis, as listed in Tables 1 and 2. The three most abundant compounds in clove volatile oil were eugenol (79.51%), CaryophylleneE (10.49%), and Caryophyllene oxide (4.40%). Terpinen-4-ol (73.53%), an organic compound of the terpene class, was the principal bioactive constituent among the three predominant components of MAEO. The second and third most common components were Camphor (10.33%) and Eucalyptol (6.09%).

**Table 1.** List of components identified in SAEO using GC–MS analysis.

| No | Retention Time | % | Components | Kovats Index | Type |
| --- | --- | --- | --- | --- | --- |
| 1 | 17.02 | 0.80 | Benzyl alcohol | 1031 | MO<sup>1</sup> |
| 2 | 31.57 | 0.09 | Cubebene<$\alpha$> | 1351 | SH<sup>2</sup> |
| 3 | 32.45 | 79.51 | Eugenol | 1359 | MO |
| 4 | 32.93 | 0.72 | Ylangene<$\alpha$> | 1375 | SH |
| 5 | 34.89 | 10.42 | CaryophylleneE | 1419 | SH |
| 6 | 36.43 | 1.33 | Humulene<α-> | 1454 | SH |
| 7 | 38.97 | 0.34 | Cadinene<δ-> | 1523 | SH |
| 8 | 39.12 | 0.34 | Eugenol acetate | 1522 | SO<sup>3</sup> |
| 9 | 39.23 | 0.31 | Calamenene<cis> | 1529 | SH |
| 10 | 41.72 | 4.40 | Caryophyllene oxide | 1583 | SO |

1: Monoterpenes Oxygenated, 2: Sesquiterpene Hydrocarbons, 3: Sesquiterpenes Oxygenated.

**Table 2.**
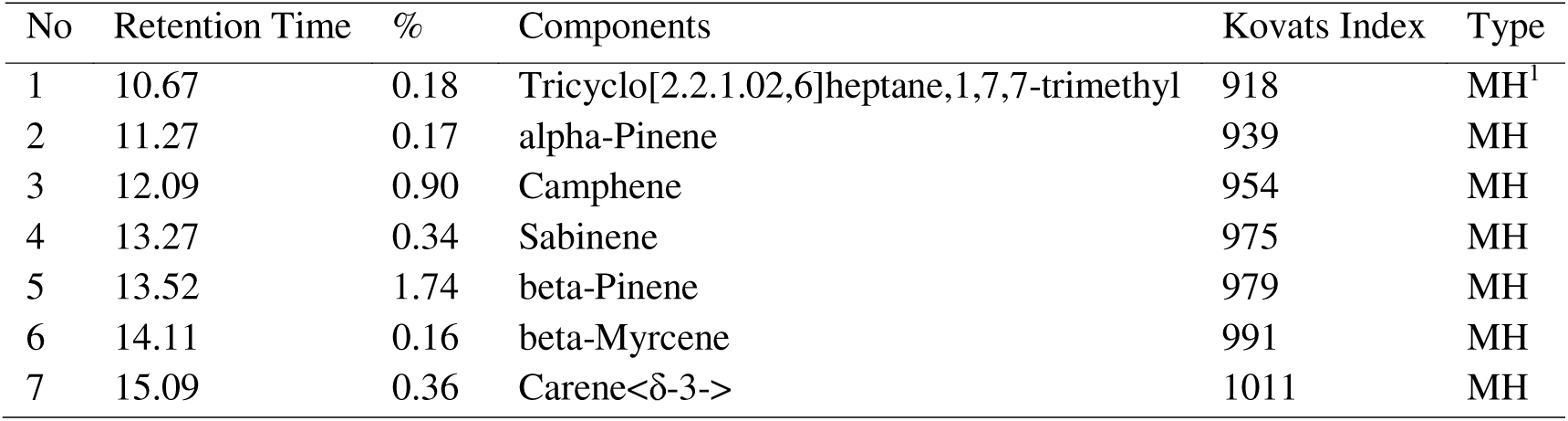
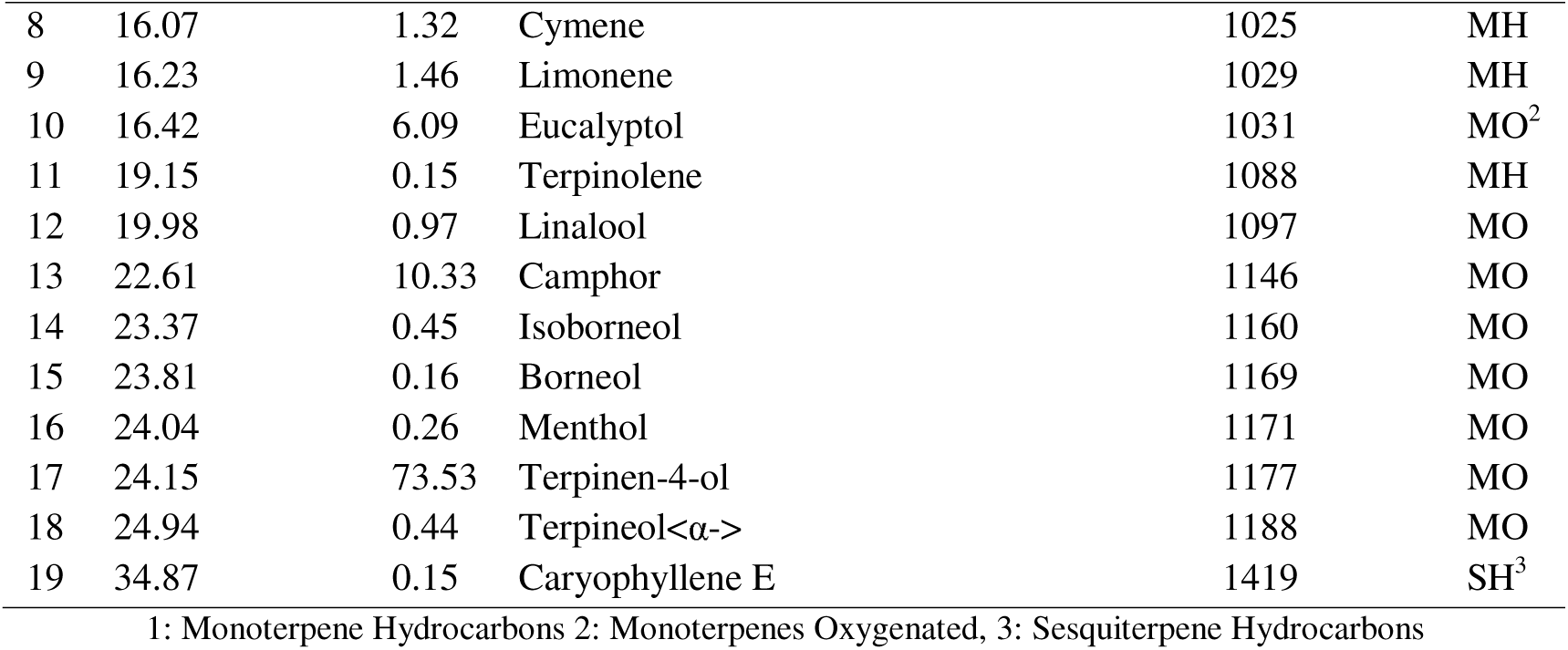
List of components identified in MAEO using GC–MS analysis.

### 3.1 Particle Size and Particle Size Distribution of the Nanoliposomes

Figure 1 depicts the DLS analyses of contributory nanoformulations. DLS profiles of the nanoliposome having 3% *S. aromaticum* EO, *M. alternifolia* EO and blank had 82.3 ± 3 nm, 102 ± 4 nm and 1.51 ± 3 µm sizes, respectively. The size distributions were calculated as 0.94 for the clove nanoliposome and 0.97 for the other samples, indicating narrow size distributions since these values were less than 1.

**Figure 1.**
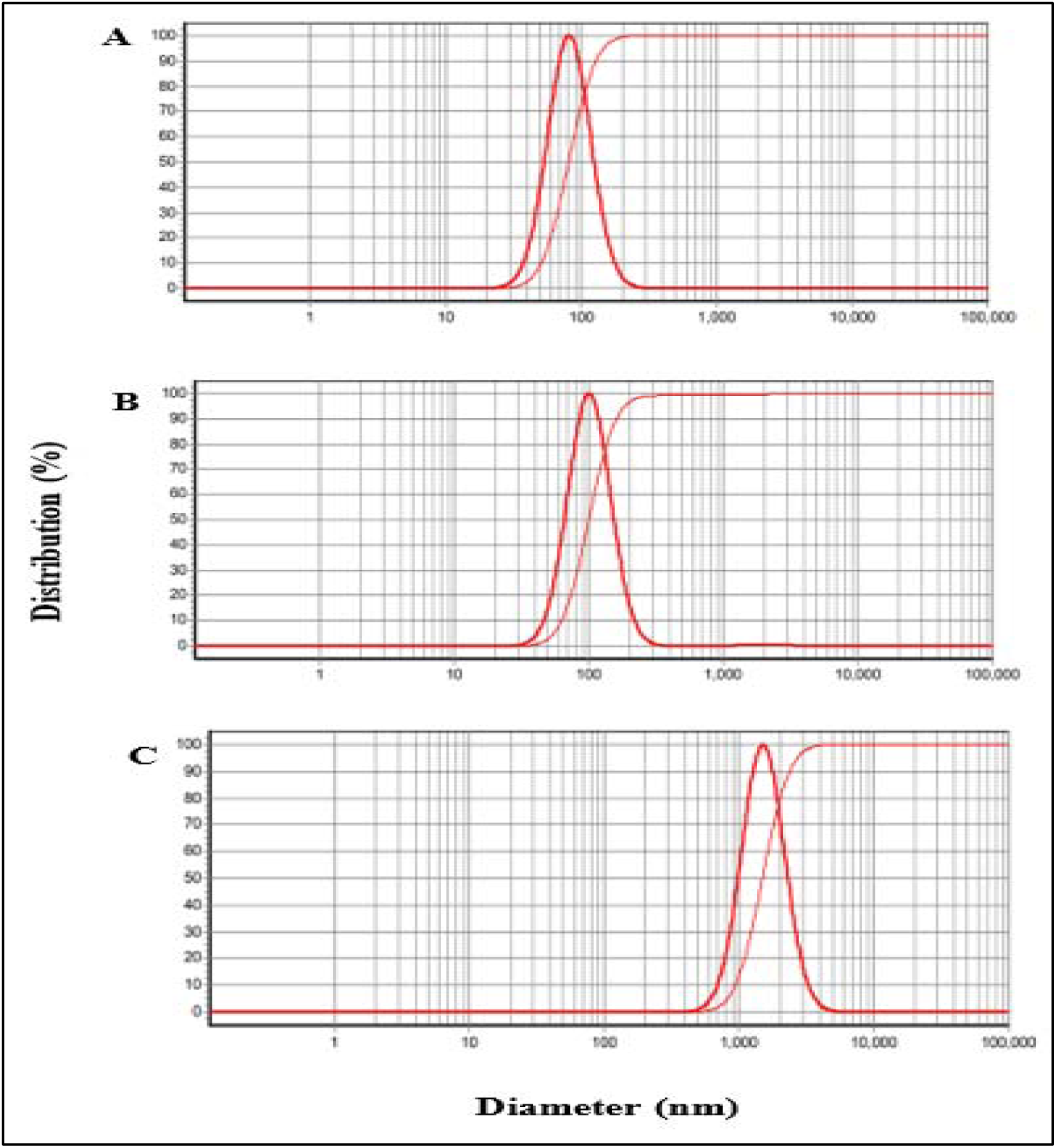
DLS analyses of A: Nanoliposomes with the 3% *S. aromaticum* EO, B: Nanoliposomes having the 3% *M. alternifolia* and C: Blank nanoliposome.

### 3.2 Physicochemical Properties of the Nanoliposomal Gel

No sedimentation, creaming, or phase separation was observed after six months of storage of the gels (LipoGel 3%, LipoGel 0.0%) at two temperatures (4^°^C and 26^°^C). Figure 2 exhibits the zeta potential profiles; the values were –38±5.8 mV, –39.1±5.1 mV and 33.3±5.7 mV, which are related to clove nanoliposome, tea tree nanoliposome, and blank nanoliposome, respectively.

**Figure 2.**
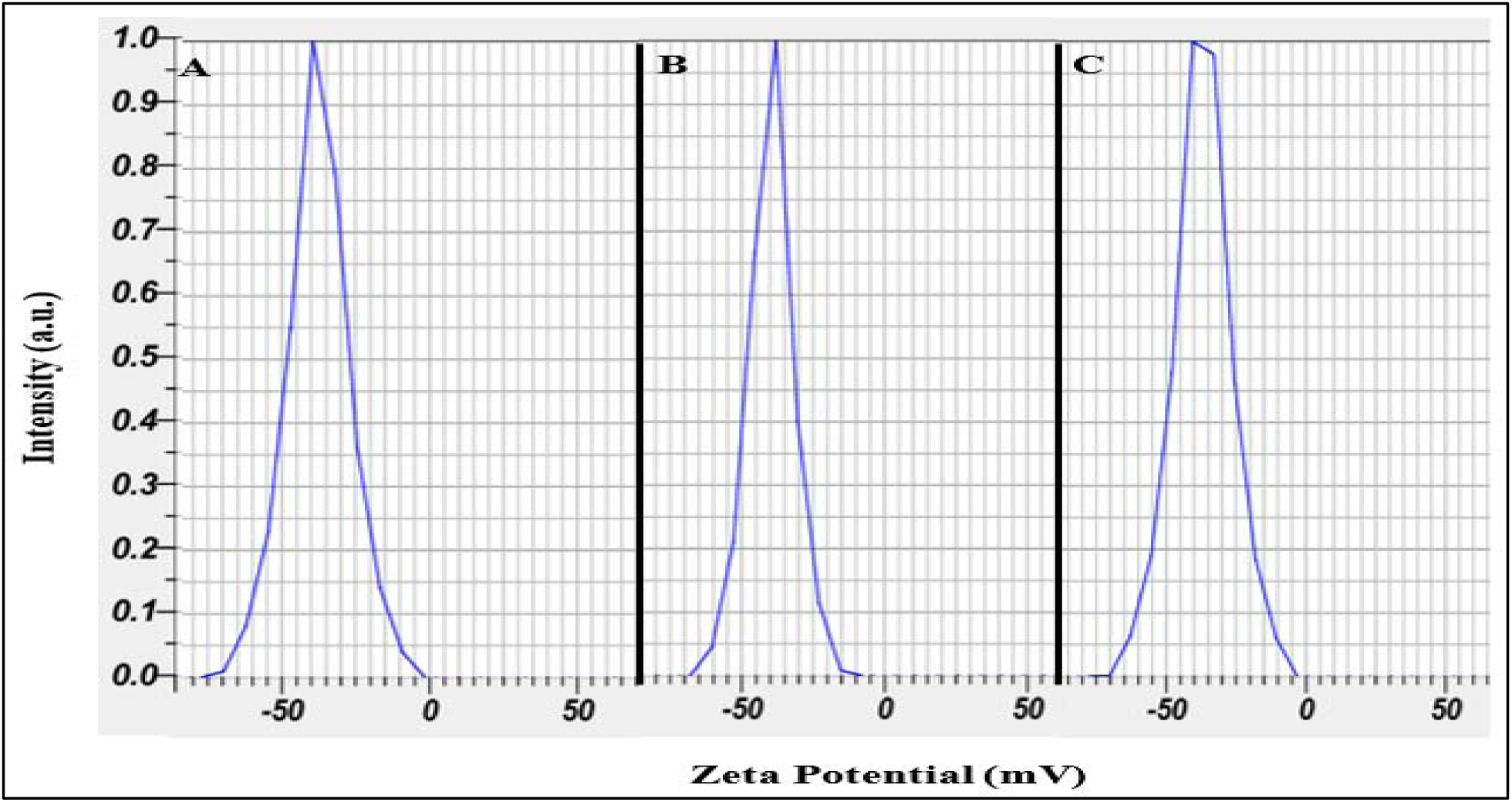
Zeta potentials of A: Nanoliposome with 3% *S. aromaticum* EO, B: Nanoliposome having 3% *M. alternifolia* EO and C: Blank nanoliposome.

Figure 3 shows the ATR-FTIR spectra of SAEO, MAEO, free nanogel, and nanogels with EO. The ATR-FTIR spectrum of clove EO showed the broad and characteristic absorption at 3515 cm^−1^ related to OH due to the phenolic compound as a major constituent in the EO, such as Eugenol, the stretching presented at 3074 and 3002 cm^−1^ related to =C-H, the spectra at 2934 and 2843cm^−1^ linked to CH_2_ vibration. The stretching bands at 1852 and 1637cm^−1^ are related to the C=O vibration; the spectra at 1607 cm, 1510, and 1457 show C=C stretching in alkene groups. The bands at 1265 and 1231cm^−1^ are attributed to C-O stretching.

**Figure 3.**
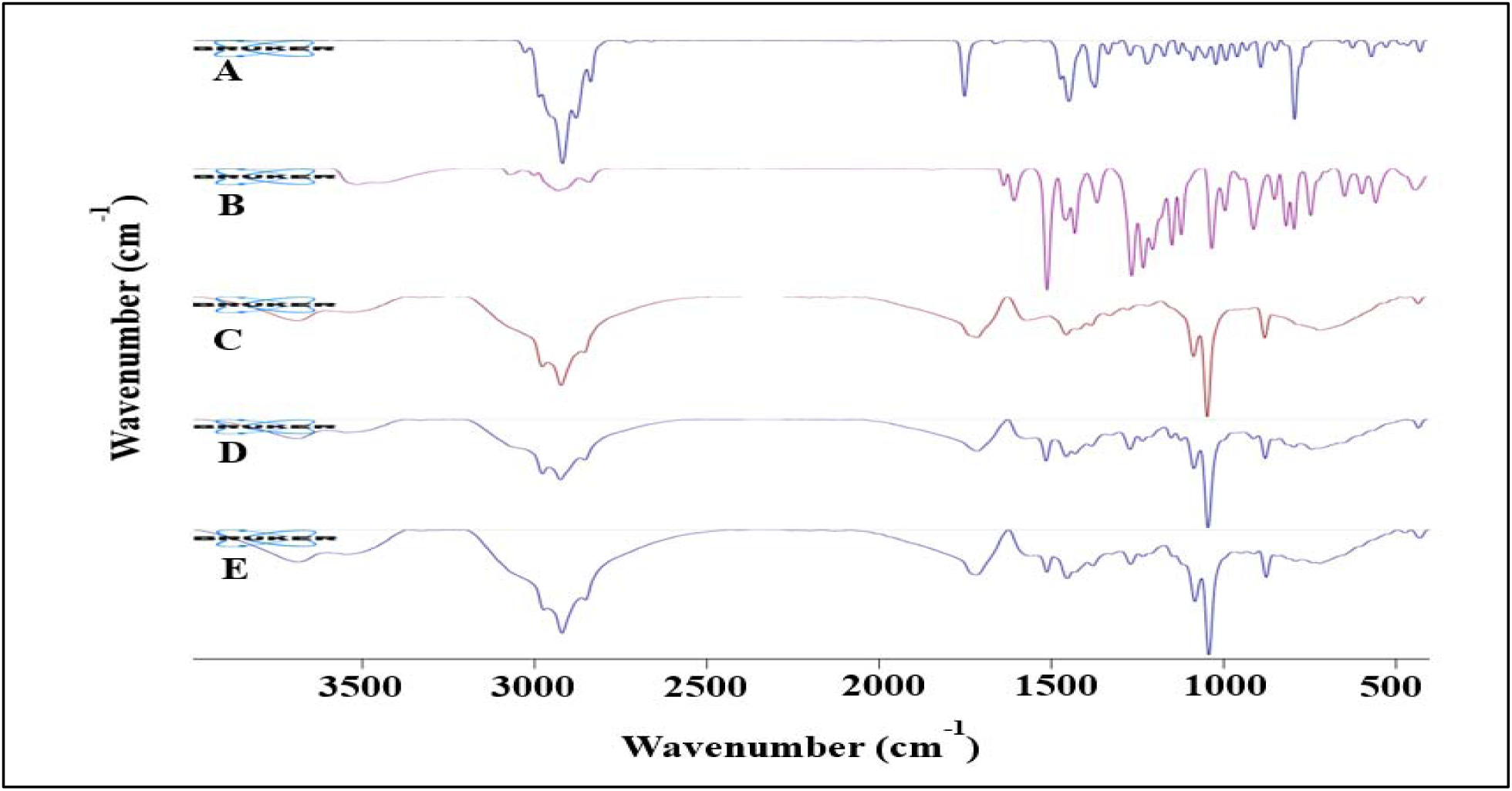
ATR-FTIR of A: *S. aromaticum* EO, B: *M. alternifolia* EO, C: Blank nanogel, D: Nanogel containing 3% *S. aromaticum* EO and E: Nanogel bearing 3% *M. alternifolia* EO.

ATR-FTIR of tea tree EO showed the C-H group of alkanes in 2983, 2916, 2877, 2834 cm^−1^; the vibration at 1746 cm^−1^ related to C=O stretch. The vibration at 1657 and 1466 cm^−1^ are attributed to the presence of C=C groups in the EO. The characteristic absorption at around 1443 cm^-1^ showed CH_2_ bending, the vibration at 1329 cm^−1^ is linked to CH_3_ group in the bioactive compound. The sharp and strong peak at 1097 cm^−1^ is attributed to C-O stretching. The strong vibration at 786 cm^-1^ is the result of C–H out of plane bending.

ATR-FTIR of blank nanogel containing the free liposome showed the hydroxyl group band in 3474 cm^−1^. The stretching bands at 2921,and 2980 cm^−1^ are related to C-H, in the alkane structures of cholesterol, lecithin, CMC and tween, the characteristic bands at 1703 cm^−1^ is related to carbonyl group in liposome. The band at 1575 cm^-1^ confirms the association of CMC with liposome through intermolecular H-bonding. The peak at around 1275 cm^−1^ is related to the phosphate group in the liposome. The sharp and strong band at 1084 and 1044 cm^-1^ are related to C-O stretching.

ATR-FTIR of the nanogel containing the clove EO liposome showed the broad and characteristic band at 3697 cm^-1^ confirming the presence of hydrogen bonding between the liposome containing clove EO and nanogel of CMC. The asymmetric stretching of CH_2_ can be observed at 2978 and 2926 and 2855 cm^-1^. The band at 1714 cm^-1^ is attributed to carbonyl group in the ester bonds of the phospholipid, the peak at around 1234 cm^-1^ related to the phosphate group in the head group of the phospholipid in liposome, these two characteristic bands demonstrated liposome structure, the spectra at 1514 is connected to C=C in aromatic ring. The vibration of carboxylic acid groups (-COOH) is responsible for a distinct peak at 1430 cm^-1^ of CMC. The sharp and strong band at 1084 and 1043 cm^-1^ corresponded to C-O.

ATR-FTIR of nanogel containing the tea tree EO liposome showed a broad band at 3542 cm^-1^ reflecting hydrogen bonding between the liposome containing tea tree EO and CMC. The asymmetric stretching of CH_2_ can be observed at 2979, 2924 and 2857 cm^-1^. The strong streching at 1714 cm^-1^ is attributed to carbonyl group in the ester bonds of the phospholipid, the peak at around 1274 cm^-1^ is related to the phosphate group in the head group of the phospholipid in liposome, that confirmed liposome structure, the stretching at 1570 is linked to C=C in aromatic ring. The vibration at 1454 cm^-1^ is related to the carboxylic acid groups in CMC. The sharp and strong band at 1084 and 1044 cm^-1^ are related to C-O stretching.

Besides, Figure 4 depicts nanogels viscosity; which was investigated at shear rates of 1 to 100 1/s. The data were fitted with the Carreau-Yasuda standard model for a non-Newtonian liquid. For all samples as the shear rate increases, the viscosity also rises.

**Figure 4.**
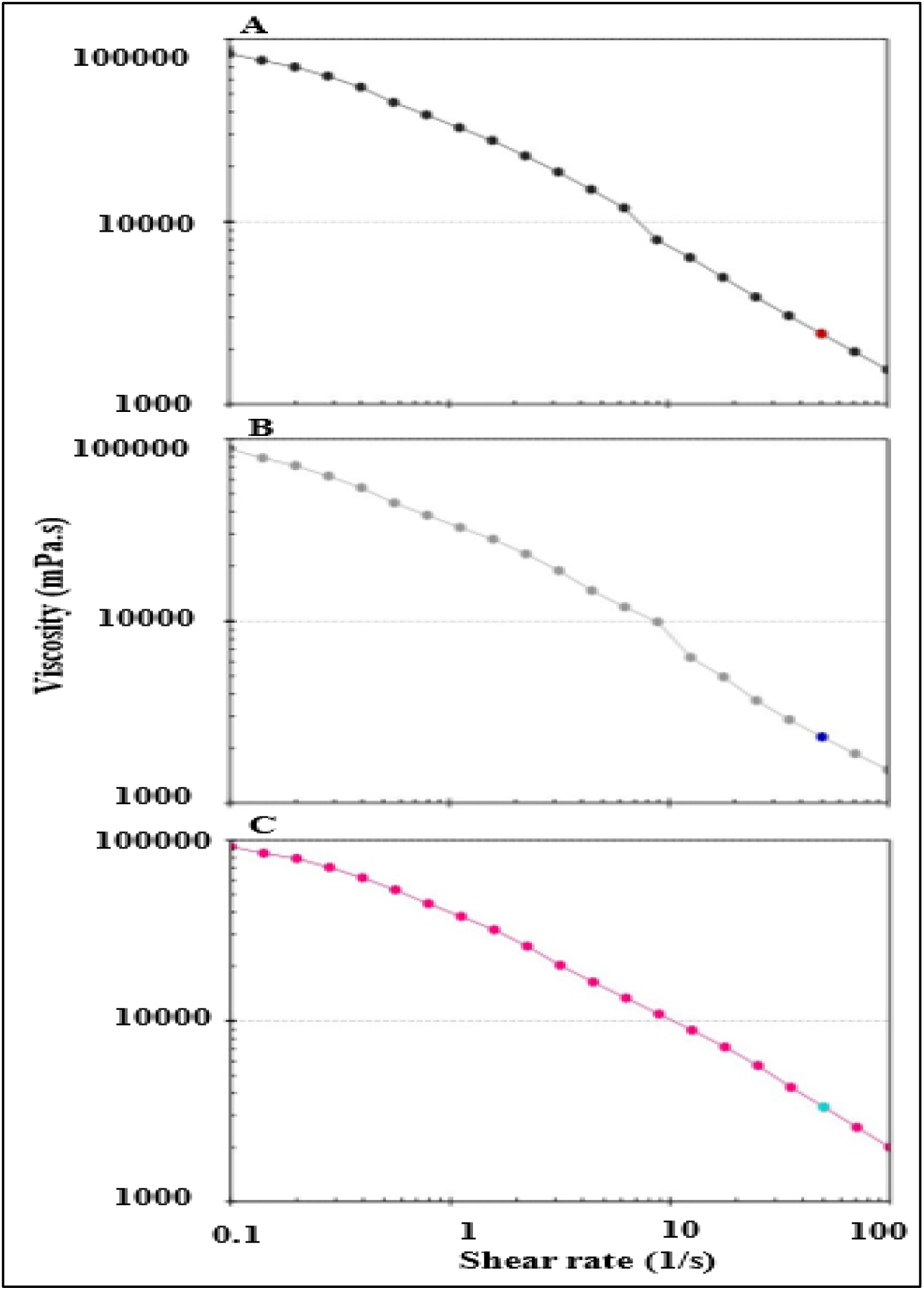
– Viscosity of A: Nanogel with 3% *S. aromaticum* EO, B: Nanogel having 3% *M. alternifolia* EO and C: Blank nanogel.

### 3.3 Repellent Effects of the Samples

Based on the data obtained from the complete protection duration test, a significant difference was observed among the different groups. The results showed that DEET (40%) had the highest protective effect, with a mean CPT of 351 ± 16 minutes. It was significantly more effective than the other samples (*P*<0.0001). Among the herbal formulations, the nanogel with clove EO demonstrated performance very close to that of DEET, with a mean protection time of 341 ± 17 minutes; the difference between clove nanogel and DEET was statistically insignificant, indicating a major increase in the efficacy of clove EO after nanoformulation. In contrast, free clove and tea tree EOs alone provided shorter protection times (45 ± 18 minutes each), which were significantly less effective than their relevant nanogels (*P*<0.0001). The tea tree nanogel also exhibited a significant improvement in protection time, with a mean of 77 ± 20 minutes compared to the free tea tree EO, but its effectiveness remained lower than that of the clove nanogel and DEET. It is noteworthy that both volatile oils formulated as nanogels showed more complete protection over time compared to the nonformulated volatile oil, which is directly attributed to the ability of the nanotechnology and encapsulation system (35). The control nanogel (blank) had the shortest protection time (34.0 ± 1.0 minutes), indicating that the observed effects were primarily due to the presence and type of EO and its nanoformulation. Statistical comparisons among samples were performed using one-way analysis of variance (One-way ANOVA) with a significance level of 0.05. The results are presented in the attached bar chart (Figure 5).

**Figure 5.**
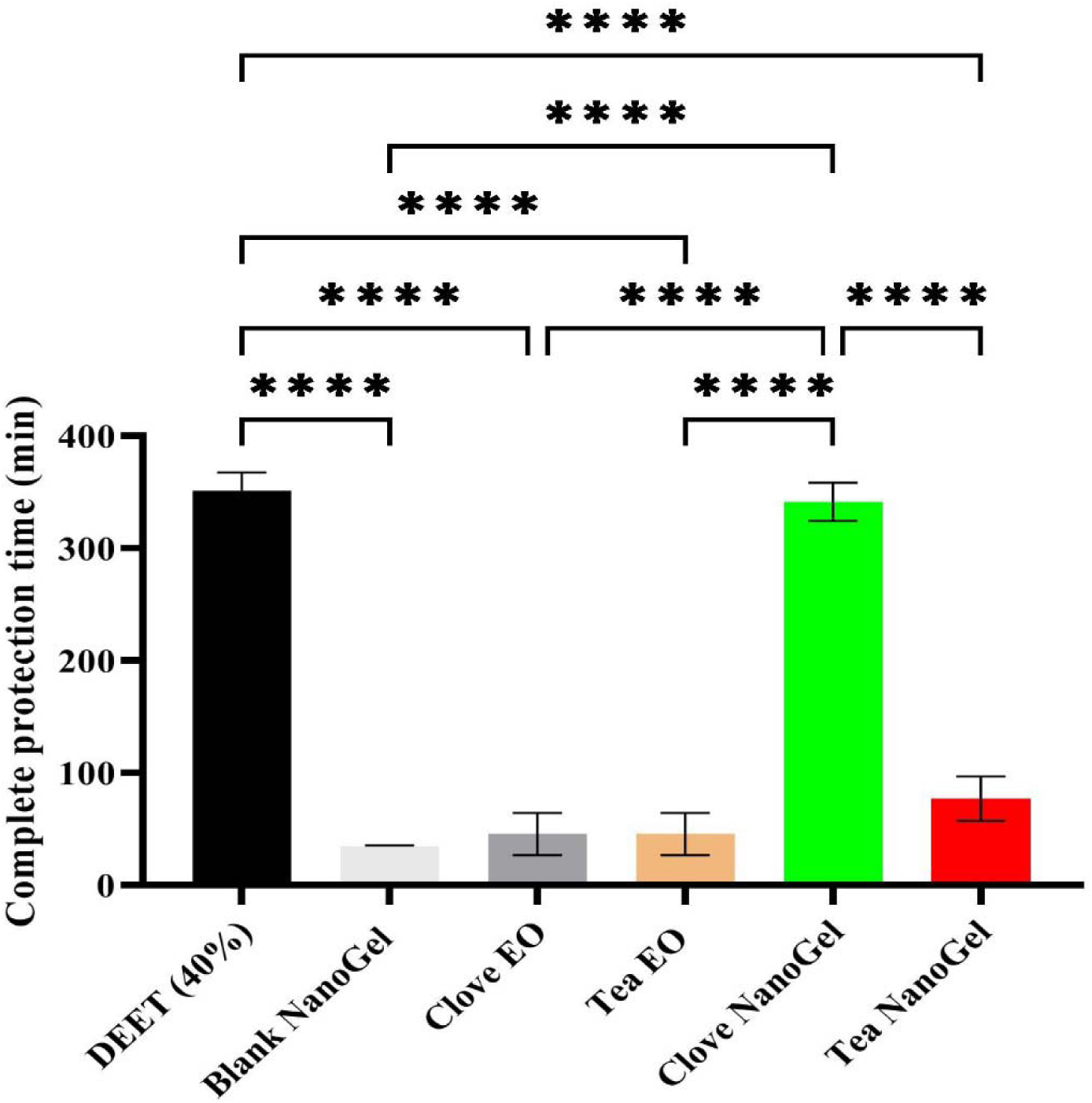
Complete protection time of the samples against *Anopheles stephensi*.

## 4. Discussion

Some mosquitoes (Diptera: Culicidae) are of great medical importance. Among them, *Anopheles stephensi* is the main vector of malaria. This mosquito has spread in many countries and in recent years has been reported as an invasive species in some countries including Djibouti, Ethiopia and Sudan (8); where it probably spread from southwest Asia. The resistance of this vector to some insecticides and larvicides as well as synthetic repellents has made it difficult to control it and eliminate malaria in some countries including Iran (12, 36, 37).

Implementing new control mechanisms and producing new formulations from natural active ingredients has thus become very important in recent years to reduce the incidence of vector-borne diseases. In order to compensate for the limitations of the use of volatile oils, the production of liposome-based nanoformulations is an effective choice (11). In this study, by adding a gelling agent (CMC), the possibility of easier and more effective topical use was also provided.

These findings could be related to the presence of the main active ingredients in clove EO, especially eugenol, which has been reported in various studies as one of the most effective compounds with insect repellent properties (22, 38, 39).

In contrast, the lower effectiveness of nanogel containing tea tree EO may be due to the difference in the nature and concentration of the active ingredients, its higher volatility, or its weaker interaction with the nanoliposomal system.

Overall, the findings of this study show that the choice of the type of volatile oil and its formulation play a decisive role in creating a repellent effect, and clove volatile oil in a nanogel substrate can be considered a more effective option to develop insect repellent products against malaria vectors. The innovation of this study is that the effectiveness of clove and tea tree volatile oils has been investigated not only individually but also in the form of nanogels.

Compared with previous studies on nanogel-based repellents against *An. stephensi*, the clove EO nanogel prepared in the present study demonstrated a remarkable repellent performance. In this study, the clove EO nanogel provided a complete protection time of 341 ± 17 minutes, which was very close to that of 40% DEET with a CPT of 351 ± 16 minutes. This finding indicates that the nanoliposomal gel formulation with clove EO could be considered a promising plant-based alternative to conventional synthetic repellents.

In a similar study, nanogels containing 2.5% EO of *Zataria multiflora*, 2.5% EO of *Elettaria cardamomum*, and 2.5% DEET showed CPT values of 600, 63 ± 15 and 242 ± 12 minutes, respectively, against *An. stephensi* (40). The CPT reported for *Z. multiflora* nanogel was considerably higher than that of the clove EO nanogel in the present study. However, the clove EO nanogel showed a much higher protection time than *E. cardamomum* nanogel and also exceeded the CPT of 2.5% DEET reported in that study. These differences may be related to the chemical composition of the essential oils, the concentration of active compounds, and the interaction of each EO with the nanogel matrix.

In another study, nanogels containing 2.5% EO of *Cinnamomum zeylanicum* and 2.5% DEET showed CPT values of 303 ± 10 and 242 ± 12 minutes, respectively, against *An. stephensi* (15). The CPT of the clove EO nanogel in the present study was higher than both the cinnamon EO nanogel and the DEET formulation reported in that study. This suggests that clove EO, particularly due to the presence of eugenol, may have stronger repellent activity or better retention in the nanoliposomal gel system.

In contrast, nanogels containing 2% EO of *Foeniculum vulgare* and 2% EO of *Mentha piperita* showed CPT values of 70 ± 6 and 120 ± 8 minutes, respectively, compared with 2% DEET with a CPT of 140 ± 8 minutes against *An. Stephensi* (41). The protection time obtained for clove EO nanogel in the present study was markedly higher than all formulations reported in that study. However, the CPT values of tea tree EO nanogel in the present study were closer to the results reported for *F. vulgare* and lower than those of *M. piperita* and 2% DEET. This comparison shows that although nanogel formulation can improve the performance of some EOs, the final repellent effect strongly depends on the type of EO and its active constituents.

Similarly, nanogels containing 2% lavender EO and 2% geranium EO showed CPT values of 90 ± 5 and 140 ± 6 minutes, respectively, while 2% DEET provided a CPT of 140 minutes against *An. stephensi* (42). The clove EO nanogel in the present study demonstrated a substantially longer CPT than lavender nanogel, geranium nanogel, and 2% DEET in that report. On the other hand, the tea tree EO nanogel in the present study showed lower or comparable protection times compared with lavender nanogel and were clearly lower than geranium nanogel and DEET. These findings further support the idea that not all plant EOs exhibit the same level of repellency after formulation in nanogel systems.

Furthermore, a nanogel containing 2.4% *Acroptilon repens* EO showed a CPT of 310 ± 45 minutes, whereas 2.4% DEET showed a CPT of 160 ± 17 minutes against *An. stephensi* (30). The CPT of the clove EO nanogel in the present study was slightly higher than that of *A. repens* EO nanogel, although the values were relatively comparable. This suggests that both clove EO and *A. repens* EO may be effective candidates for the development of plant-based repellent formulations. However, the clove EO nanogel showed a more favorable result compared with the DEET concentration used in that study.

In a study with a similar design, nanogels containing 3% carvacrol, cinnamaldehyde, and thymol were evaluated against *Aedes aegypti* and compared with 40% DEET. The CPT values were 200 ± 34, 250 ± 34, and 240 minutes, respectively, while DEET showed a CPT of 190 ± 17 minutes (43). Although this study was conducted on *Ae. aegypti* rather than *An. stephensi*, the clove EO nanogel in the present study showed a higher CPT than all three nanogel formulations and DEET reported in that study. This may indicate the strong repellent potential of clove EO in nanoliposomal gel form, although direct comparison should be made cautiously because of differences in mosquito species, formulation composition, and test conditions.

In addition, a study on nanoliposomal hydrogel containing 3% *Pelargonium* EO, nonformulated 3% *Pelargonium* EO, and 3% DEET against *An. stephensi* reported CPT values of 216 ± 25, 148

± 23, and 311 ± 37 minutes, respectively (35). The clove EO nanogel in the present study showed a longer CPT than both formulated and nonformulated *Pelargonium* EO and was also slightly higher than the reported CPT of 3% DEET. This comparison highlights the potential advantage of clove EO nanogel over some other plant-based formulations.

Another study showed that solid nanoparticles containing 1% *Zataria multiflora* EO had a CPT of 93 ± 5 minutes, while 1% nonformulated EO showed a CPT of 29 ± 2 minutes against *An. stephensi* (17). The CPT of the clove EO nanogel in the present study was much higher than both nanoparticle-based and nonformulated *Zataria multiflora* EO. However, the CPT values of tea tree EO nanogel in the present study were closer to the nanoparticle formulation of *Z. multiflora*. These findings suggest that formulation type, EO concentration, and chemical profile are critical factors influencing the duration of protection.

Overall, comparisons with previous studies indicated that the clove EO nanogel developed in the present study exhibited one of the highest CPT values among several plant-based nanogel or nanoparticle formulations evaluated against *An. stephensi*. Its CPT was close to that of 40% DEET and higher than many previously reported EO-based formulations.

Finally, it is an inescapable paradigm that despite the unequivocable pace of research in the field of controlling infectious diseases in recent decades, the challenge of combating vector-borne zoonotic, and specifically anthroponotic, maladies remain a formidable task to overcome (44–46).

## Conclusions

These findings highlight the potential of nanoliposomal based nanogel repellents, particularly those containing clove, as effective and safe botanical alternatives to conventional repellents, with promising implications for integrated vector management strategies. It could thus be postulated that the nanogel of clove EO can be considered as a natural and environmentally-friendly repellent for further investigation against mosquitoes.

## Data Availability

The data generated or analyzed during this study are available from the corresponding author on reasonable request.

## Ethical Approval

The study has been ethically approved by the Ethics Committee of the Shiraz University of Medical Sciences IR.SUMS.SCHEANUT.REC.1404.02. The process of the mosquito repellent assays was fully described to the candidates. As a result, they volunteered to participate in this research. Moreover, all methods in the current study were performed according to the WHO relevant guidelines and national and international regulations.

## Consent

Informed consent was obtained from all participants.

## Acknowledgments

The authors acknowledge the valuable assistance of the Vice Chancellor for Research and Technology, Shiraz University of Medical Sciences (SUMS) and the excellent logistical support provided by the Department of Medical Nanotechnology, Fasa University of Medical Sciences (FUMS). This paper was part of our Master’s thesis in Biology and Control of Disease Vectors by Ms. R. Shahheidari, with a grant (Number 31676) awarded to her supervisor, Dr. M.D. Moemenbellah-Fard under the Code of Ethics: IR.SUMS.SCHEANUT.REC.1404.02. The authors are also very grateful to all their respected colleagues at FUMS and SUMS and several colleagues, Dr. M. Shahriari-Namadi, and other members of the technical staff for their valuable assistance in facilitating the desired outcome of this project.

## Authors’ contributions

R Shahheidari, MD Moemenbellah-Fard, M Osanloo collectively endorsed the proposal, performed the experimental works, contributed to data gathering, and revised the original draft. A Paksa, re-analyzed data and statistical methods. AH Roozitalab and M Fakhraei, collaborated in breeding mosquitos and conduct repellency tests. E Zarenezhad, performed, analyzed, and wrote the ATR-FTIR spectroscopy analysis. All authors contributed to the writing and final verification of ultimate draft.

